# Thermal sensitivity and seasonal change in the gut microbiome of a desert ant, *Cephalotes rohweri*

**DOI:** 10.1101/2021.12.28.474384

**Authors:** Marshall S. McMunn, Asher I. Hudson, Ash T. Zemenick, Monika Egerer, Stacy M. Philpott, Rachel L. Vannette

## Abstract

Microorganisms within ectotherms must withstand the variable body temperatures of their hosts. Shifts in host body temperature resulting from climate change have the potential to shape ectotherm microbiome composition. Microbiome compositional changes occurring in response to temperature in nature have not been frequently examined, restricting our ability to predict microbe-mediated ectotherm responses to climate change. In a set of field-based observations, we characterized gut bacterial communities and thermal exposure across a population of desert arboreal ants (*Cephalotes rohweri*). In a paired growth chamber experiment, we exposed ant colonies to variable temperature regimes differing by 5 ^°^C for three months. We found that the abundance and composition of ant-associated bacteria were sensitive to elevated temperatures in both field and laboratory experiments. We observed a subset of taxa that responded similarly to temperature in the experimental and observational study, suggesting a role of seasonal temperature and local temperature differences amongst nests in shaping microbiomes within the ant population. Bacterial mutualists in the genus *Cephalotococcus* (Opitutales: Opitutaceae) were especially sensitive to change in temperature—decreasing in abundance in naturally warm summer nests and warm growth chambers. We also report the discovery of a member of the Candidate Phlya Radiation (Phylum: Gracilibacteria), a suspected epibiont, found in low abundance within the guts of this ant species.

## 1. Introduction

Temperature change is a ubiquitous challenge faced by living organisms. Warming produces conformational changes in proteins, alters enzyme efficiencies, and can eventually lead to protein denaturation (Vogt, Woell and Argos 1997). A large majority of multicellular life on earth, by total biomass and diversity, is ectothermic (Bar-On, Phillips and Milo 2018), relying on external heat to regulate body temperature. As ectotherm body temperature fluctuates, so does the temperature experienced by associated microbiomes. Across diverse animal taxa, microbiome composition and abundance is sensitive to thermal change, however, examples linking natural variation in temperature to changes in microbiome composition remain rare (Moeller 2004; Dunbar *et al*. 2007; Lemoine, Engl and Kaltenpoth 2020; Onyango *et al*. 2020). The thermal biology of microbiomes may be an important mechanism affecting ectotherm response to climate change because of the many host traits that are linked to microbiome composition and abundance, including pathogen resistance, nutrient acquisition, and reproduction (Sepulveda and Moeller 2020).

Temperature variation across seasons, days, and habitat plays a large role in determining terrestrial ectotherm species distributions and strongly shapes the evolution of thermal tolerance of organisms (Huey, Berrigan and Miles 2001). Many observations of ectotherm thermal tolerance across geographic gradients and life histories have accumulated to shape the field of thermal biology. As an example, optimal temperatures of ectotherms tend to be higher at low latitudes where temperatures are warmer, while thermal tolerance breadth (range of suitable temperatures) is narrower in these same areas due to more stable temperatures across seasons (Stevens 1989; Addo-Bediako, Chown and Gaston 2000). Worldwide, there is a pattern among terrestrial ectotherms of thermal maximums being constrained and varying less over geographic gradients than species thermal minimums (Snyder and Weathers 1975). The degree to which these and other global patterns of adaptation in response to variation in temperature are also reflected in thermal responses of microbiomes within ectotherms is largely unknown. Additional examples of ectotherm microbiome response in nature may aid in testing this generalization.

Ectotherm responses to temperatures of different extremes or duration may also guide expectations of microbiome thermal sensitivity (Iltis *et al*. 2021). Acclimation, a gradual improvement in ability to tolerate temperature deviations from optimum following exposure, is very common among ectotherms (Gaston *et al*. 2009). Extreme temperatures (e.g. heat shock) can be harmful even if the exposure is brief, while smaller deviations from optimum temperature may be tolerated for long periods of time without severe consequence (Heath 1963; Kingsolver and Gomulkiewicz 2003). Studies that examine microbiome response to warming suggest that adaptive acclimation can occur with gradual exposure (Moghadam *et al*. 2018), irreversible damage can occur from extreme exposure (Kikuchi *et al*. 2016), and gradual recovery can follow moderate exposure (Heyworth, Smee and Ferrari 2020). We do not have a clear picture of how these responses may align or interact with host biology at present moment, and thus more examples of natural responses are needed.

Changes in the abundance of microbial strains can be driven directly by microbial temperature sensitivities or through a wide array of indirect mechanisms that can be difficult to experimentally separate (Corbin *et al*. 2016). Indirect mechanisms may include temperature dependent changes in host physiology that alter the microbial environment (e.g. gut pH), complex interactions amongst microbes due to asymmetric temperature responses, or microbial evolution and co-evolution within novel environmental contexts (Sepulveda and Moeller 2020).

Much of the knowledge of microbiome response to temperature has come from growth chamber experiments, which offer precise control over thermal exposure (Russell and Moran 2006; Dunbar *et al*. 2007; Fan and Wernegreen 2013) or lab-based microbial growth and colonization assays (Hammer, Le and Moran 2021). However, relatively few studies have investigated whether temperature alters microbiome composition in nature or if changes in nature are temporary. The difficulty in characterizing *in situ* microbiome temperature sensitivities is due to both the challenge of characterizing thermal exposure in free-living animals and the number of correlated biotic factors that change along with temperature (e.g. seasonal diet shifts) (Maurice *et al*. 2015).

To further improve predictions of the fate of ectotherms in a changing climate, it will be critical to examine microbiome response to thermal stress and also validate it in a natural setting. With this aim we characterized microbiome composition and abundance of a desert ant, *Cephalotes rohweri* (Hymenoptera: Formicidae) by pairing a growth chamber experiment with an observational study of seasonal and microclimate nest temperatures. We aimed to answer the following questions: 1) Do nest microclimate and seasonal temperature correlate with microbiome composition and abundance in the field? 2) Are correlations between temperature and composition in the field replicated in growth chambers? and 3) What microbial taxa are sensitive to environmental temperature, and can we explain any of the variation in responses across the natural population?

## 2 Methods

### 2.1 Summary

We studied the effects of temperature on the gut bacteria of *Cephalotes rohweri* by comparing the composition and abundance of gut bacteria across a population of ant colonies in two seasons (summer – late September and winter – late February). We then exposed collected whole ant colonies to daily temperature patterns differing by 5^°^C in a growth chamber experiment (Figure 1). Our study utilized 16S amplicon sequencing and 16S quantitative PCR from ant midguts and ilea to characterize gut microbiome composition. All analyses were performed in R (R version 4.1.1) (R Core Team 2020).

**Figure 1:**
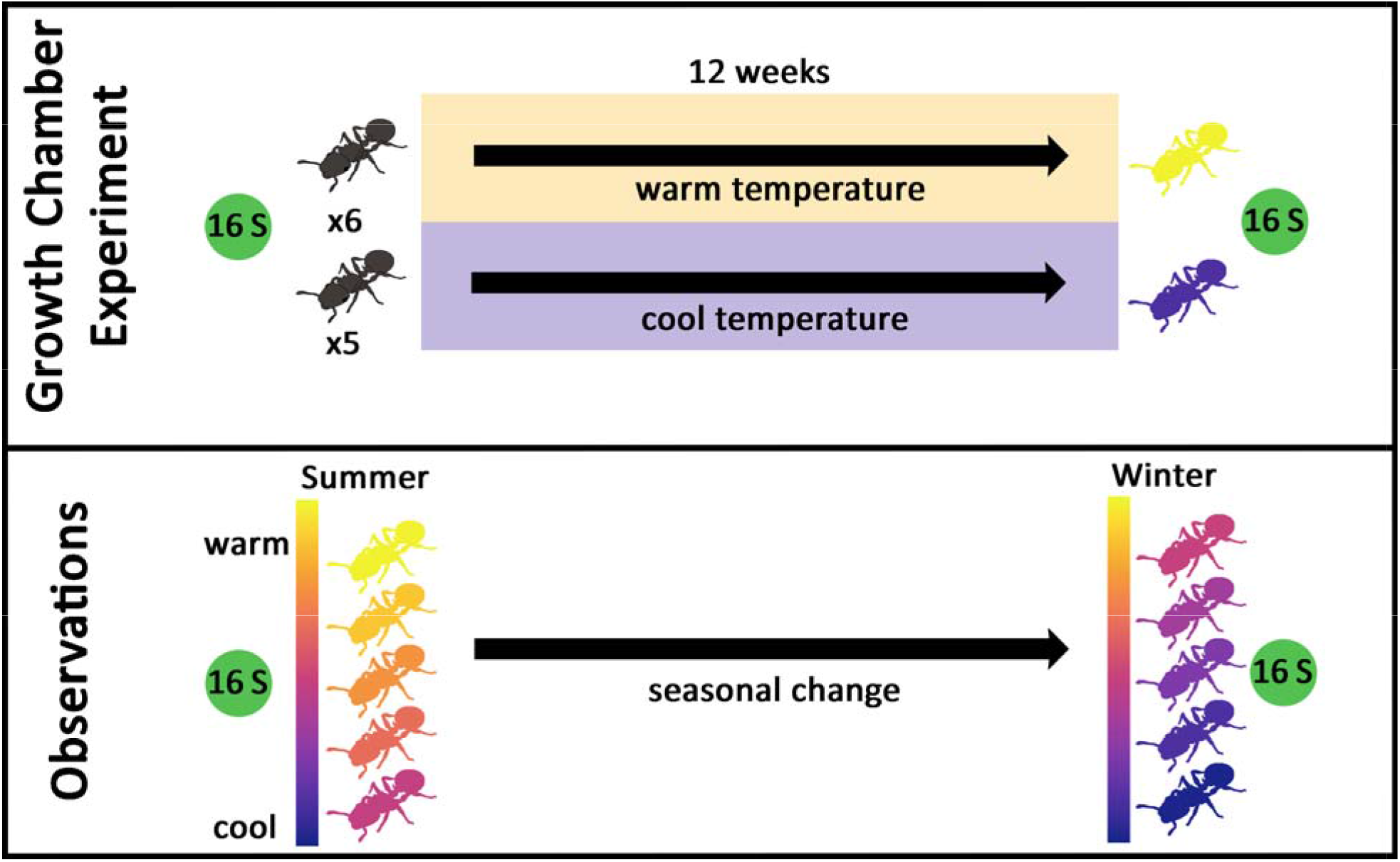
Outline of experimental and observational approaches. Top panel - *Cephalotes rohweri* ant colonies were moved into growth chambers programmed to warm (+1^°^C) and cool (−4^°^C) temperatures relative to observed hourly average nest temperatures in the field in the summer. Microbiome composition and abundance was assessed with 16S amplicon sequencing and qPCR. Bottom panel - In a complementary set of observations, we characterized naturally occurring nest temperatures within both the summer (September 2018) and winter (February 2019) and naturally occurring bacterial composition. We also characterized seasonal bacterial compositional change as we resampled the same colonies the following winter.

### 2.2 Study species

*Cephalotes rohweri* live in the dead branches of several tree and shrub species in the Sonoran Desert. Nests are occupied up to several years at a time, with a small proportion of workers leaving the nest at any time for foraging. Colonies of *C. rohweri* form a reproductive unit and consist of 3 castes, including a single queen, a minor caste (workers), and a major caste (soldiers) (Powell and Peretz 2021). These individuals are spread between one to several nests in a single tree, with minor workers being the most abundant caste and responsible for most of the resource acquisition. As arboreal ants living in a desert, *C. rohweri* are exposed to extreme summer temperatures and moderate winter temperatures inside their poorly insulated nests (Figure 2).

**Figure 2.**
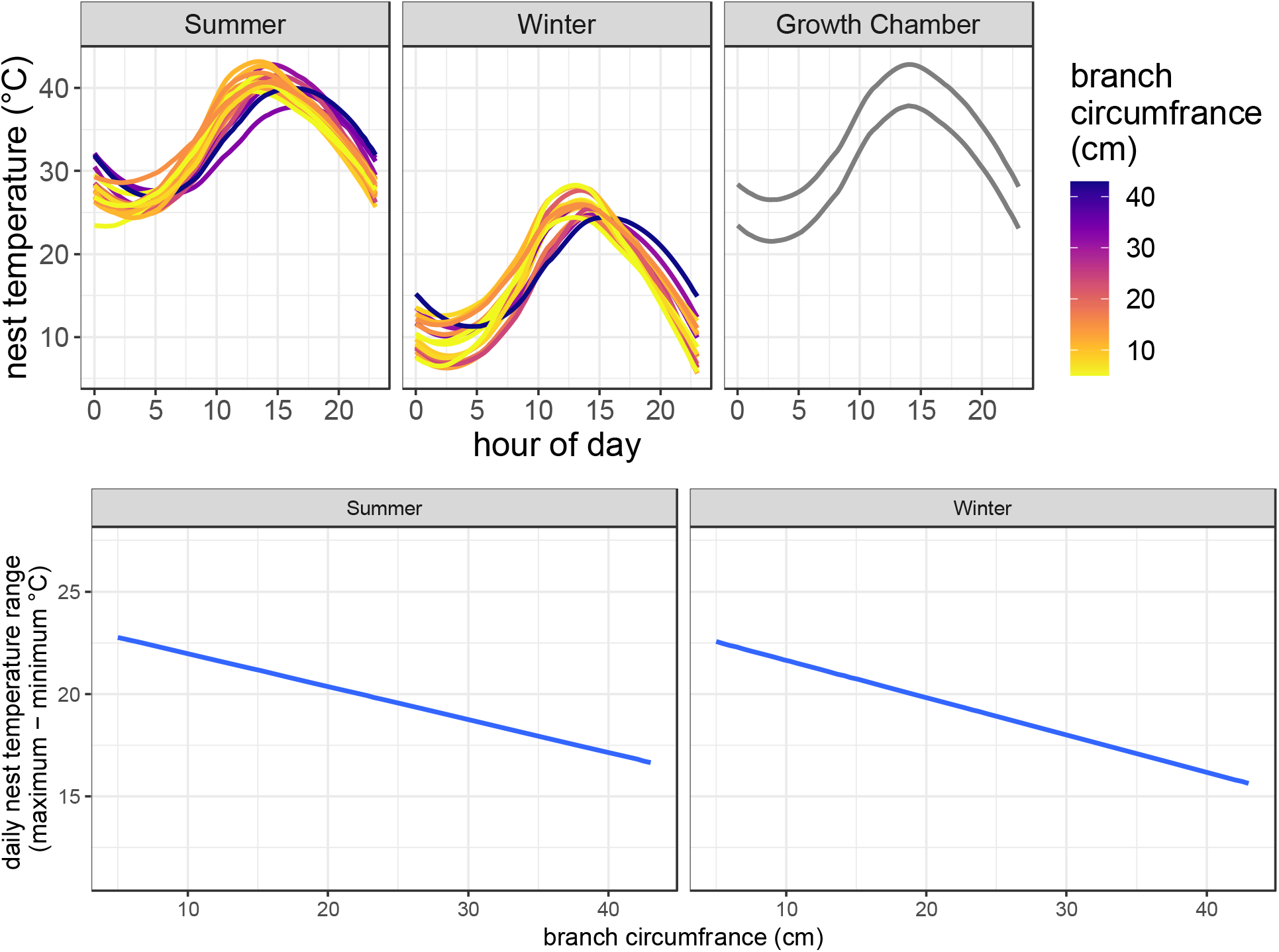
Top panels: smoothing splines of median hourly temperature colored by nesting branch diameter across growth chamber experiments, summer nest temperatures collections, and winter collections. Growth chamber treatments fall within natural summer ranges but represent approximately -4^°^C and +1^°^C each hour for “cool” and “warm” respectively. Bottom panels: summer and winter nest temperature range in ^°^C. Both summer (R^2^ = 0.22, p = 0.016) and winter (R^2^ = 0.25, p = 0.023) with larger branch circumferences have smaller temperature ranges.

Abundant bacterial mutualists within the midgut and ileum of *C. rohweri* upgrade otherwise inaccessible nitrogen compounds (urea, uric acid, and xanthine) to produce amino acids (Hu *et al*. 2018). Through metagenomic sequencing, metabolic pathway reconstruction, and controlled experiments, Hu et al. (2018) demonstrated that *C. rohweri* realize a very efficient nitrogen economy, with bacteria enabling ant survival on an otherwise poor diet largely consisting of plant exudations, bird excrement, pollen, and likely fungal spores. Ants of the entire genus *Cephalotes* have low trophic positions compared to other ants, rarely if ever taking animal prey (Russel *et al*. 2009). All *Cephalotes* species investigated thus far, including 17 of 119 total species in the genus (Powell, Price and Kronauer 2020), possess microbiomes with broadly similar composition and functional capabilities in terms of core bacterial taxa (Hu *et al*. 2018) suggesting a long co-evolutionary history between ants and at least some bacterial lineages. *Cephalotes rohweri* is the most northerly distributed species of this diverse neotropical ant genus.

### 2.2 Study site

We collected *C. rohweri* nests and individuals from branches of foothills palo verde trees *(Parkinsonia microphylla*) within the Tucson Mountain Range (32.242^°^N, 111.093^°^W) in the Arizona Upland region of the Sonoran Desert (southeastern Arizona, USA). The habitat is typical of a Sonoran thornscrub with abundant Saguaro cacti (*Carnegiea gigantea*), foothills palo verde, and a diversity of other native Cactaceae, small shrubs, and flowering annual plants. The area experiences frequent hard frosts in the winter, summer daytime temperatures regularly reaching 45^°^C, and annual monsoon rains.

### 2.4 Minor worker collections – microclimate and season

In the summer (early September 2018) and the following winter (late February 2019), we collected minor workers using aspirators foraging within 0.5 m from 19 identified nest entrances. Ants were kept in a cooler in the shade for a maximum of 6 hours until they were frozen each afternoon upon return from the field.

Interior nest temperature measurements corresponding to September 2018 (summer observations) and February 2019 (winter observations) were taken using a J-type thermocouple and a HOBO UX100 thermocouple datalogger. The thermocouple wire was inserted 0.5-2 cm past the nest entrance and recorded interior nest temperature every minute for between 36 and 96 h. Summary statistics from nest temperature data independently used in analysis include mean, median, maximum, minimum, 95% quantile, and 80% quantile temperatures. Our choice of these variables was based on the desire to quantify the temperatures that microbes were exposed to for the longest duration, but also investigate the effects of short-lasting regular exposure to more extreme high temperatures. We used a linear regression to test for an effect of nest branch exterior diameter on the daily range of temperatures within the nest.

### 2.5 Growth chamber experiment

To test the effects of temperature on ant gut bacterial composition and abundance we performed a 3-month growth chamber experiment, comparing a warm and cool treatment. The two temperature treatments were based on field measurements of daily temperature trajectories observed within nests but differed by 5^°^C from each other (Figure 2). Hourly temperatures were calculated from 7 months (March 2017 - September 2017) of temperature readings within dead tree branches at the collection site using iButtons (DS1922L, Maxim Integrated Products, San Jose, CA, USA) (Figure 2). When we examined summer nest interior measurements of greater accuracy (September 2018), we found our treatments were well within the bounds of natural variation observed in nests, approximately 1^°^C above average hourly summer temperatures (warm) and 4^°^C below average hourly summer temperatures (cool) (Figure 2).

We collected 11 colonies in September 2017 by removing all potential nest-containing branches (those with dead wood) from trees in which we observed foraging *C. rohweri*. Nest entrances were blocked with Play-Doh (Hasbro, Inc. Pawtucket, RI) for up to 72 h prior to manual nest dissection. All individual ants from a single tree regardless of caste or developmental stage were placed into an artificial nest made from a 12.7 cm x 17.8 cm picture frame with a 5mm entrance hole drilled into the side. The clear plexiglass was covered with red-tinted tape. Colonies were ordered by total number of workers and alternately assigned to cool (5 colonies) and warm (6 colonies) temperature treatments and placed in their respective growth chambers with a 12-h light, 12-h dark regime setting.

Ants were fed ad libitum outside the artificial nest with wicking feeders replaced twice each week with DI water, NaCl solution, and urea solution. The foraging area for each nest consisted of a 32 cm x 20 cm x 14 cm plastic container with its walls coated with fluon (Insect-a-slip, Bioquip, Rancho Dominguez, CA). Outside of feedings, the top was covered with plastic wrap and rubber banded in place to prevent escape. Small dishes were used for 50% honey solution (absorbed onto a folded and mounted KimWipe), bee pollen, and 1 frozen cockroach nymph. Between 2-5 live ants were dissected and sequenced from each colony three times: at setup, after 6 weeks of exposure, and after 3 months of exposure.

### 2.6 Ant gut dissections, DNA extractions, and sequencing

We dissected minor workers from growth chamber experiments and field collections under sterile conditions in Dubesco’s phosphate buffered solution, removing the midgut and ileum from each ant. Samples were bulked to obtain adequate concentrations of DNA (1-5 minor workers in 2017 and 2 minor workers in 2018/2019). Forceps were flame-sterilized with ethanol between ant digestive tract dissections, and sterile buffer and petri dishes were used for each dissection. Control samples consisted of extracted DNA from forceps that were dipped into the dissection liquid following an ant dissection and were sequenced for each round of dissections.

DNA was extracted from the bulked midguts and ilea using a modified Powersoil (Qiagen, Hilden, Germany) extraction protocol, including an additional tissue disruption step followed by an overnight Proteinase K soak (Rubin *et al*. 2014). To increase DNA concentration, the final elution step was performed with half the recommended volume of the final elution buffer (50uL solution C6). DNA concentration was measured using a Nanodrop 1000 spectrophotometer prior to sample submission to confirm successful DNA extraction (ThermoScientific, Waltham, MA, USA).

Extracted DNA from samples and controls for each round of dissections were sequenced by the Centre for Comparative Genomics and Evolutionary Bioinformatics Integrated Microbiome Resource at Dalhousie University. Primers were selected to align with the Earth Microbiome project, specifically the V4 subregion of 16S SSU rRNA - 515fB (GTGYCAGCMGCCGCGGTAA) and 806rB (GGACTACNVGGGTWTCTAAT). DNA was amplified using High fidelity Phusion polymerase. Amplified DNA was sequenced using an Illumina MiSeq, producing 291 bp paired end reads.

We also performed quantitative PCR (qPCR) in triplicate for each sample to estimate the per ant 16S copy number. We used the same primers above (V4 subregion of 16S SSU rRNA), SSOAdvanced Universal SYBR Green Supermix kits (BioRad Laboratories, Inc., Hercules, CA, USA), and a QFX96 thermocycler (BioRad) to quantify DNA concentration over 35 PCR cycles (94^°^C for 45s, 50C for 60s, 72C for 90s) with a 10-minute extension at 72^°^C. We averaged the 3 Ct values for each sample, and calculated estimated copy number based on the difference between Ct values of the sample and a calibrated between plate standard. Samples were distributed across 3 plates randomly with respect to treatment groups and collection times. Our between plate standard consisted of 5uL aliquots from 15 DNA extractions bulked across treatment groups, sampling times, and extractions. Read count was estimated using this standard of known 16S copy number (Tkacz, Hortala and Poole 2018). The qPCR standard was produced by isolating the 291 bp 16S band from a post-PCR gel, performing a DNA extraction using the procedure above, calculating 16S double-stranded DNA concentration using a Qubit DNA assay (Invitrogen, Waltham, MA), and then converting DNA concentration (ng/uL) to copy number using the molecular mass of the 291 bp fragment.

### 2.7 Amplicon sequence variant, taxonomic, and phylogenetic assignments

Illumina 16S reads were processed using the DADA2 workflow to obtain amplicon sequence variants (ASV) including filtering, dereplication, inference of sequence variants, merger of paired-end reads, and chimera removal (Callahan *et al*. 2016). The DADA2 pipeline was applied independently to each Illumina run (growth chamber experiment, summer observations, winter observations). These 3 runs were separated to accommodate possible variation in error rates and to allow likely contaminants to be identified within each dataset prior to merging. Likely contaminant ASV’s were identified using the package *decontam*, applying a 50% prevalence threshold, removing ASV’s more prevalent in controls than true samples, and ASVs occurring in less than 3 samples. Taxonomy was assigned to ASVs using DADA2 assignTaxonomy function, in combination with the ribosomal database project naïve Bayesian classifier algorithm and using the Silva training dataset (v138) (Wang *et al*. 2007). We used the phanghorn R package to build a de novo phylogenetic tree of the 3 merged datasets using the neighbor-joining method (Schliep 2011).

### 2.8 Statistical analyses of response to temperature

We assessed the response of ant gut bacterial communities to temperature between warm and cool growth chamber treatments, across natural microclimatic gradients within summer and winter samples, and seasonally between summer and winter samples (Figure 1).

We multiplied our estimated qPCR per-ant 16S read abundance (from here on qPCR weighted abundance) for each sample by relative abundance of each ASV. This approach provided a quantitative estimate of ASV abundance within each sample (Jian *et al*. 2020). We tested for compositional dissimilarity in abundance of ASVs across samples and homogeneity in beta dispersion across treatment groups and seasons using betadisper and adonis (package *vegan* 2.5) (Oksanen *et al*. 2020). These analyses were performed for all sampling dates in the experiment separately (0 months, 1.5 months, and 3 months in growth chamber) and between summer and winter observations.

We investigated differences in qPCR-weighted abundance using DESeq2 (Love, Huber and Anders 2014) at the ASV, genus, family, and order level using a single response of growth chamber treatment, season, or a single selected observed temperature summary statistic (for the two microclimate analyses). Each set of microclimate comparisons (summer and winter) was investigated by performing single predictor variable PERMANOVA’s with each contending temperature summary statistic (mean, median, 80^th^ quantile, and 95^th^ quantile, minimum, and maximum temperature) and comparing output. The two seasons are dramatically different in temperature regimes and following model comparison, the two summary statistics explaining the greatest variation (95^th^ percentile for summer microclimate, and 80^th^ percentile temperature for winter microclimate) were used in all subsequent analyses (DESeq2, UNIFRAC) for these respective datasets. For summer temperature, 95^th^ quantile temperature may be capturing the negative effects of sustained elevated daytime temperature. For winter, the opposite may be occurring, with microbial growth being slowed during the cooler hours of the day.

To test for phylogenetic signal in thermal response of bacterial clades, we analyzed both weighted and unweighted UNIFRAC distances comparing warm and cool growth chamber ants after 3 months of exposure, ants collected in winter versus summer, and according to microclimatic variation within both summer and winter. Both unweighted and weighted UNIFRAC distances were used to determine if our predictor variables (season, treatment, or microclimate) change the presence and absence of closely related bacterial taxa in synchrony (unweighted), but also whether they shift the abundance of closely related bacterial taxa in synchrony (weighted). As above, we used the 95^th^ percentile nest temperature as a predictor variable in the summer and the 80^th^ percentile nest temperature for the winter.

To estimate temperature effect on the overall bacterial abundance in ant guts, we used linear models along microclimate gradients (predictor variables – 95^th^ percentile and 80^th^ percentile temperature for summer and winter respectively), ANOVA (between final timepoint lab-based warm and cool treatments), and paired t-tests (seasonal observations) to test for the difference in estimated qPCR 16S log read count per ant. We averaged all 3 qPCR technical replicates before converting to an estimate of per ant log 16S read abundance. For growth chamber experiments, we report change in per-ant 16S read abundance from initial measurements for each colony after 3 months of exposure.

Pearson correlations of the log-fold change in response to temperature for all ASVs were used to determine if taxa responded similarly across growth chamber experiments, between seasons, and along microclimatic gradients. For microclimate, per-degree log-fold change was used from the output of DESeq2.

## 3. Results

### 3.1 Summary

We identified 173 unique ASVs from 3,035,597 reads, representing 7 orders of bacteria, including Betaproteobacteriales (64 ASVs), Opitutales (29), Xanthomonadales (29), Rhizobiales (22), Pseudomonadales (18), Flavobacteriales (9), and JGI_0000069P22 (phylum Gracilibacteria) (2). This closely resembles previous studies of microbiome composition in this species.

We found that bacterial symbionts in the guts of *C. rohweri* are sensitive to temperature changes in growth chambers, across seasons, and across summer and winter microclimatic gradients. In particular, ASVs in the family Opitutaceae were reduced in abundance in the nests that experienced the highest temperatures during the summer (95^th^ percentile temperature).

Arboreal ants are more vulnerable with regards to temperature fluctuations compared to ground-nesting ants. We found that nest temperature range depended on branch size with larger branches being less susceptible to daily temperature changes in both the summer (F_1,19_ = 6.93, R^2^ = 0.22, p = 0.016) and winter (F_1,15_ = 6.37, R^2^ = 0.25, p = 0.023) (Figure 2).

### 3.2 Field sampling: seasonal change

There were strong seasonal differences in bacterial species composition within the ant population (F_1,34_ =13.77, R^2^=0.28, p<0.001) (Figure 3, Figure 4). Seasonal changes in bacterial abundance were detected in 6 orders, 5 families, 2 genera, and 42 ASVs significantly differed between summer and winter samples (Supplemental Table 1 FDR<0.05, Figure 5)

**Figure 3.**
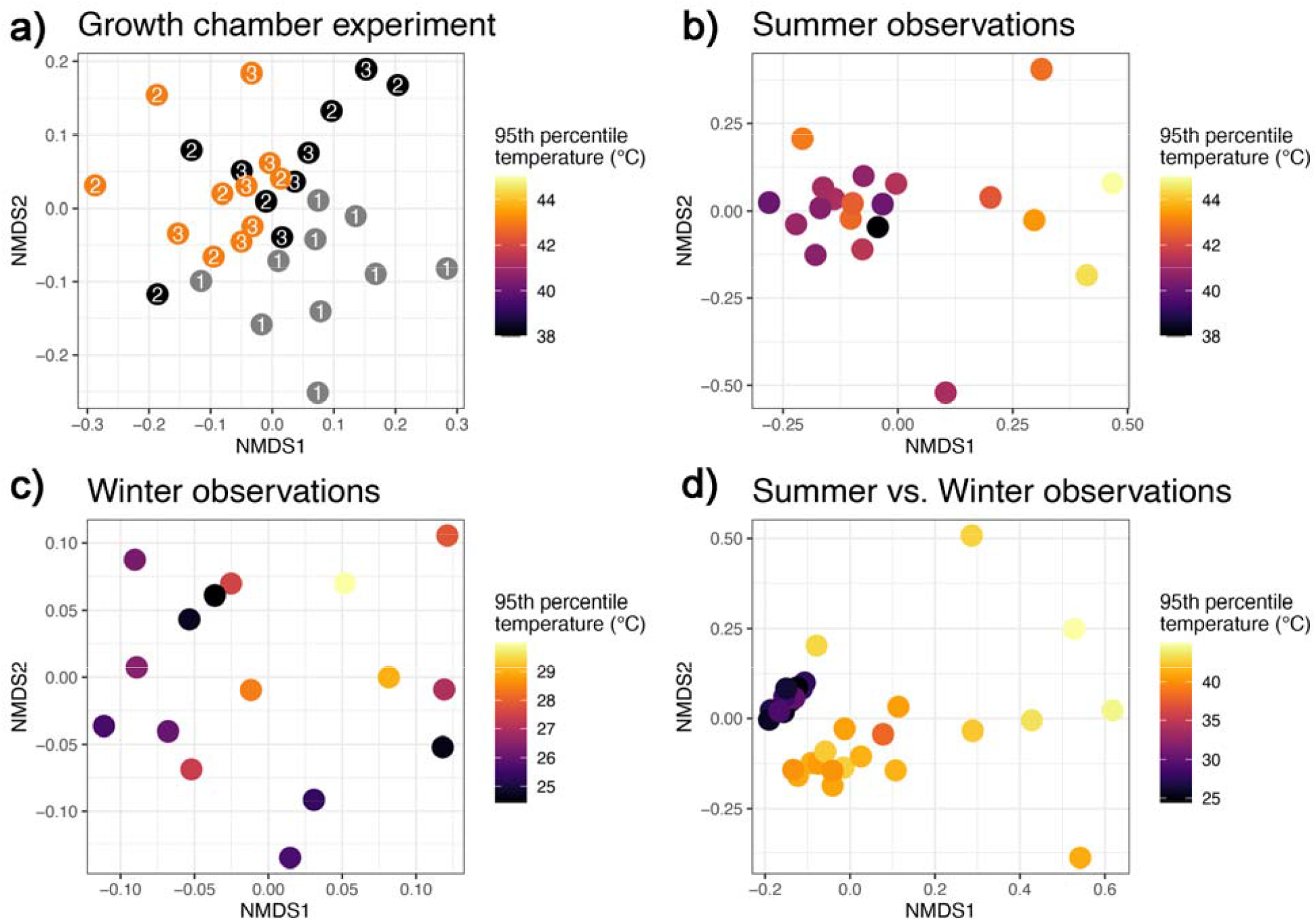
NMDS of qPCR weighted bacterial community composition collected from *C. rohweri* ant guts with each point representing a separate sample of bulked ants from a separate nest. a) growth chamber experiment. Color corresponds to treatment, with initial conditions (grey) and warm (orange) and cool (black) treatments. Labels of points correspond to initial (1), 6-week (2), and 3-month (3) sampling periods. b) summer 2018 field collections colored by nest temperature and c) 2019 winter field collections with each point representing bulked ants within distinct colonies. Color scales are the same for panel a and b – with 95^th^ percentile temperature in degrees C for the growth chamber treatments and nest temperature measurements.

**Figure 4.**
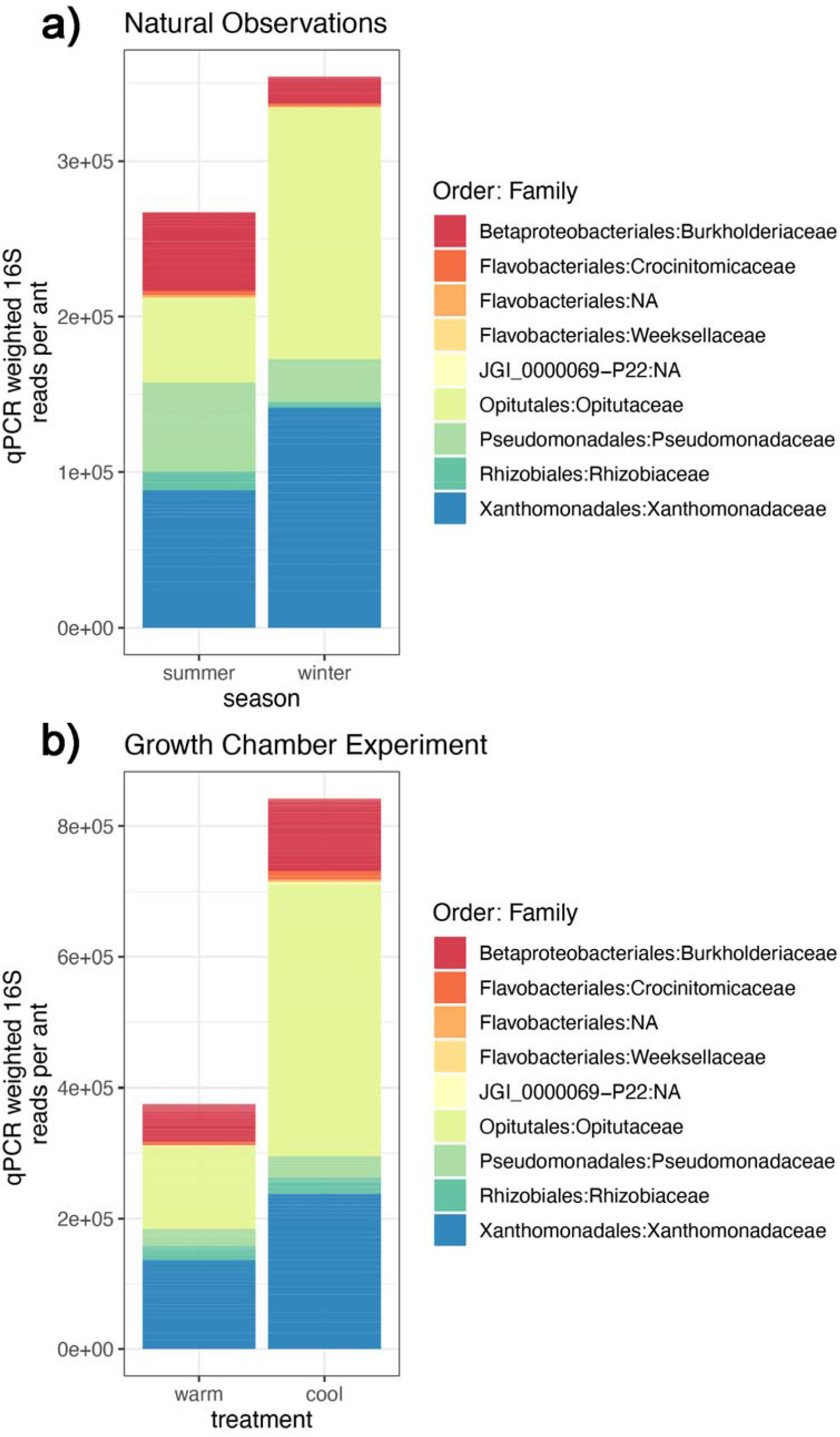
Stacked barplots of qPCR weighted abundance of all ASVs color coded by family and order a) 2017 temperature treatments and b) summer and winter field collections. There are more bacteria in both the cooler temperature treatment and the winter nests in nature and broadly (total height), and compositional shifts were similar between the paired treatments and observations (correlated log-fold changes of ASVs in Figure 5).

**Figure 5.**
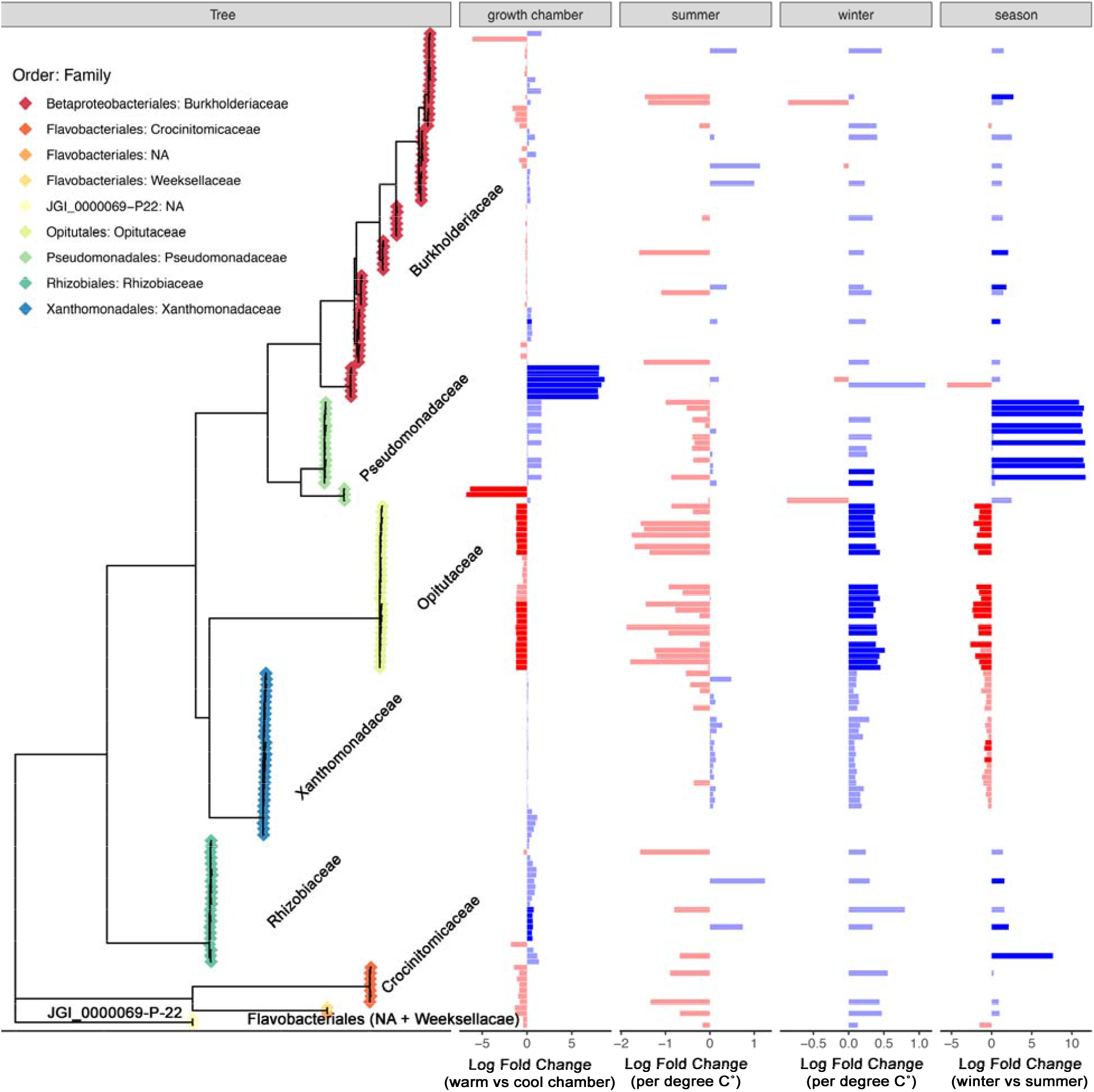
log-fold change in qPCR weighted relative abundance of all ASVs across temperature ASVs demonstrating significant response to temperature or treatment (resulting from DEseq2 analyses) are shaded with 100% opacity. ASV’s unchanged in 16S qPCR weighted abundance are shaded with 70% opacity. Red bars represent a negative response to increased temperature, while blue bars represent a positive response to increased temperature. 1) phylogenetic tree of all ASV’s colored by taxonomic family. 2) log-fold change number of 16S copies in response to growth chamber treatments. 3) per degree log-fold change number of 16S copies in response to summer microclimate. 4) per degree log-fold change number of 16S copies in response to winter microclimate. 5) log-fold change number of 16S copies between winter and summer (blue means greater in summer, red means greater in winter). Members of the Opitutaceae were particularly sensitive to temperature change and were lower in abundance in the warm growth chamber, lower in the summer, when aggregated lower in warm summer nests compared to cool summer nests, and higher in abundance in warm winter nests than cool winter nests.

Weighted UNIFRAC phylogenetic distance between samples differed significantly between seasons (F_1,35_=94.09, R^2^ = 0.729, p<0.0001), suggesting that more related bacteria respond similarly in terms of seasonal change in abundance. Neither unweighted UNIFRAC distance nor alpha diversity differed by season, suggesting that within host ASV gains or losses between seasons are not phylogenetically clustered. Samples in the winter were more homogenous than the same colonies measured in the summer (betadisper, F_1,34_=73.64, p<0.001). We observed 35% more total 16S read copies per ant in winter colonies compared to summer colonies (paired t-test, t=2.19, df=15, p = 0.045).

### 3.3 Variation with microclimate

Exposure of individual ant nests to extreme summer temperatures, as measured by 95^th^ percentile nest temperature, was correlated with variation in bacterial composition and diversity among nests. The 95^th^ percentile temperature predicted variation in ant gut bacterial composition (16S qPCR weighted) (PERMANOVA, F_1,18_=2.73, R^2^ = 0.13, p=0.015) (Fig. 3b). No other temperature explanatory variables (mean, median, 80^th^ percentile temperature) explained a significant amount of variation in summer community composition. Change in qPCR-weighted abundance in response to 95^th^ percentile nest temperature in the summer was significant (via DESeq2 analyses) in one aggregated family (*Opitutaceae*), one genus (*Cephaloticoccus*), and 4 ASVs (all ASVs *Cephaloticoccus, Opitutaceae*) (Figure 5, Supplemental Table 1). Both Weighted (presence/absence) and unweighted (abundance) UNIFRAC phylogenetic distance varied with 95^th^ percentile nest temperature in the summer (weighted, F_1,18_ = 3.51, R^2^ = 0.16, p = 0.016) (unweighted, F_1,18_ = 4.03, R^2^ = 0.18, p = 0.034). This suggests that the impact of high summer temperatures depended on phylogeny, both in terms of their presence and absence (unweighted) and abundance (weighted). As nest temperature decreased in summer nests we observed higher bacterial alpha diversity (t=-2.82, R^2^ = 0.26, p = 0.011).

Winter microclimate also influenced community composition, but only when using 80^th^ percentile temperature as the predictor variable (F_1,14_ = 3.32, R^2^ = 0.19, p = 0.011). No other temperature explanatory variables (mean, median, 95^th^ percentile temperature) explained a significant amount of variation in winter community composition (Fig. 3c). Change in bacterial relative abundance in response to 80^th^ percentile nest temperature in the winter was significant in one family and one order (Xanthomonadales, Xanthomonadaceae) and 23 ASVs – Supplemental table 1) (Figure 5). Neither alpha diversity nor weighted and unweighted UNIFRAC phylogenetic distance between samples varied with winter nest temperature.

There were no significant differences in total 16S read copy number qPCR measurements along microclimatic gradients within the summer (t=-0.01, p=0.99) or winter (t=1.42, p=0.17) nests according to temperature predictors.

### 3.4 Growth chamber experiment

We found that bacterial community composition was sensitive to temperature treatments after 3 months of exposure (adonis, R^2^ = 0.27, F_1,10_ = 3.43, p=0.036), but no difference was detected 6 weeks after exposure (adonis, R^2^= 0.15, F_1,9_=1.47, p=0.24), suggesting gradual change resulting from temperature exposure in growth chambers (Fig. 3a). Warm nests had lower among-nest variance in species composition than did the control nests after 6 weeks of temperature treatments (betadisper, F_1,8_=7.79, p<0.025), while we found no difference in between treatments after 3 months.

A total of 36 ASVs significantly responded to growth chamber treatments—including ASVs in the genus *Cephaloticoccus* which decreased in abundance at warm temperatures (Supplemental Table 1, Figure 5). At higher taxonomic levels, no aggregated orders or families responded significantly to growth chamber treatments, but the aggregated genus *Cephaloticoccus* decreased in relative abundance in the warm temperature treatment.

We observed a large decrease over the course of the experiment (initial – 3 month) in perant log 16S read abundance in warm temperature treatments relative to the reduced temperature treatment (F_1,8_ = 9.94, p<0.013), suggesting that bacterial abundance in ant guts was lower following 3 months at warmer temperatures.

Weighted UNIFRAC phylogenetic distance between samples differed significantly between growth chamber treatments (F_1,9_=4.38, R^2^ = 0.33, p = 0.024). Unweighted UNIFRAC distance between samples (p>0.05), as well as bacterial alpha diversity, did not differ between warm and cool treatments, suggesting that we did not see within host gain or loss of related ASVs as was the case across seasons.

### 3.5 Growth chamber vs. naturally observed thermal responses

Bacterial ASVs that were rare or abundant in hot summer nests showed similar changes in abundance between cool and warm growth chambers through a positive correlation between log-fold change across growth chambers treatments and per degree log fold change across the population of nests in the summer (R = 0.30, t = 2.84, p = 0.005) (Figure 5). We saw a similar pattern when comparing summer microclimate response with growth chamber treatments, finding a positive correlation between log-fold change across growth chambers treatments and seasonal shifts in the ant’s naturally occurring microbiome (R = 0.19, t = 2.53, p = 0.012). We found no correlation between winter temperature responses and growth chamber treatments.

### 3.6 Presence of members of the phyla Gracilibacteria

Two ASVs were annotated as phylum Gracilibacteria (order JGI_0000069-P22). These ASVs were detected in multiple colonies and at least one occurred in each of the 3 years of sampling. While this ASV was detected in low abundance (maximum of 214 reads in a sample), we detected presence of the order-level taxon JGI_0000069-P22 in 46 of 69 16S sequencing samples and none of the 6 control samples sequenced.

## 4. Discussion

*Cephalotes rohweri* gut bacterial composition and abundance are sensitive to naturally occurring variation in temperature due to seasonal change and nest microclimate. Our major results indicate that 1) mutualists decrease in abundance in summer nests, but seasonally recover by late winter within colonies and 2) winter bacterial composition shows less variance than summer composition, which when considered alongside decreased summer 16S read abundance, suggests that summer ant colonies may be experiencing dysbiosis.

Our results demonstrated that bacteria in the genus *Cephaloticoccus* (Opitutales: Opitutaceae), which enhance nitrogen metabolism in this species of ant (Hu *et al*. 2018), were particularly sensitive to changes in temperature in growth chambers between seasons, and in warmer nests within the natural population during the summer. We found members of *Cephaloticoccus* at reduced abundance in warm naturally occurring summer nests and cold naturally occurring winter nests, suggesting a narrow thermal range (sensitive to both hot and cold extremes) compared to other bacteria within these ants. Differences observed between ant nests in the summer did not persist through the following winter, suggesting seasonal recovery from temperature exposure within colonies in nature. Seasonal changes in microbiome composition have been noted in several other ectotherm taxa, with varying effects on host phenotype (Ferguson *et al*. 2018; Liu, Lei and Chen 2019; Zhao *et al*. 2021).

In our growth chamber experiment, we found reduced overall bacterial abundance in the warm treatment compared to the cool treatment. While we demonstrate here that the bacterial abundance and composition of this ant is sensitive to temperature, we caution that functional assays are required to assess if such losses affect host fitness. Direct assays of microbiome efficiency are necessary to confirm shifts in microbiome-based nutrient acquisition in the summer as either bacterial efficiency or host ability to utilize nutrients could change in tandem with abundance and composition of the ant microbiome (Hu *et al*. 2018).

Thermal sensitivity occurs not only across seasons but according to microhabitat, with warm and cool nests collected from within a very small area (800 m x 500 m) varying significantly in microbiome composition. These findings indicate that change in temperature across very fine scales of time and space could play an important role in shaping host microbiome composition, and warrant study in additional systems of animals. Colony temperature measurements in the field explain variation in microbiome composition, even given potential sources of error in temperature measurements over a short period of time (1-3 days), the potential for between nest worker movement within colonies, and the many other factors that vary between ant colonies (e.g. food access or age/size). Given the small spatial scale at which we detected differences between ant colonies in the summer and then convergence toward a more similar winter microbiome composition across the population, we suggest that the microbiome of *C. rohweri* may shift in composition regularly and reversibly in response to natural temperature fluctuations.

*Cephalotes rohweri* may be able to buffer against permanent loss of microbial strains at the colony level due to social sharing of microbiota – one of several potential mechanisms through which microbial associations may be recovered following thermal stress. New workers, present to some extent in nests across seasons, are inoculated by nestmates through oral-anal trophallaxis (Lanan *et al*. 2017). Assuming microbes were not lost by every colony member simultaneously and entirely, this mechanism of transmission could reestablish heat sensitive taxa despite loss in individual colony members. This could be resolved in future studies that did not pool individuals within colonies.

Two of the main bacterial groups that were demonstrated to respond to thermal change include Opitutales and Xanthomonadales. These groups have been previously characterized in other *Cephalotes* species using metagenome assembled genomes. Both are notable for their abundance, strain diversity, and roles in nitrogen metabolism, as both encode urease accessory proteins in *C. rohweri* (Hu *et al*. 2018).Interestingly, only in the species *C. rohweri* do *Cephaloticoccus* genomes lack a full complement of urease accessory proteins. These proteins are instead found only within a member of the Xanthomonadales, a clade that in all other *Cephalotes* microbiomes lacks this urease functionality all together (Hu *et al*. 2018). We do not currently understand the detailed co-evolutionary history of the *C. rohweri* bacterial symbionts, but this combined gain and loss of functionality in these two temperature sensitive microbial clades only within *C. rohweri*, suggest the possibility of recent co-evolution – potentially including horizontal gene transfer from *Cephaloticoccus* to Xanthomonadales.

While the results presented here suggest that key members of the microbiome are sensitive to thermal stress, we cannot address the functional consequences for ant hosts. Transplanting microbiomes between individuals exposed to novel conditions into unexposed individuals would represent a more direct test of the effects of abiotically-induced compositional shifts in the microbiome. Such experiments have been conducted in *Drosophila melanogaster* and aphids and have demonstrated that the effects of past exposure to temperature on the microbiome can be inherited and have consequences for the host phenotype (Moghadam *et al*. 2018; Heyworth, Smee and Ferrari 2020). Transplant experiments have linked temperature-driven microbiome changes to improved temperature tolerance, but beneficial or detrimental microbiome compositional shifts may occur in a much wider array of animals (Sepulveda and Moeller 2020; Iltis *et al*. 2021). This type of experiment may be difficult to conduct in *Cephalotes* spp., as microbiomes are believed to be largely isolated from external contamination after formation of the proventriculus in newly enclosed adults, leaving only a short window in which to perform experimental microbiome transplants (Lanan *et al*. 2017).

Microbe-based acclimation and microbe-based susceptibility surely both occur in nature, and host physiology, diet, and habitat likely matter a great deal in determining the direction of this outcome (Sepulveda and Moeller 2020). Identifying general characteristics that allow for predictions of host microbiome responses to changes in abiotic conditions, and in particular temperature, will be key in understanding how a host will fare under climate change. We suggest that *C. rohweri* in particular has several life history characteristics that may result in acclimation rather than susceptibility. These traits include functional redundancy across closely related bacterial ASVs (Figure 5, this paper (Hu *et al*. 2018) within individual microbiomes and additional redundancy of microbial taxa across the super organism. Reproduction within ants occurs at the colony level, and in this species by a single queen within that colony, and social sharing of microbes is common in this species (Lanan *et al*. 2017). In addition to redundancy of individuals, colonies frequently consist of multiple sizes of branches, each containing a nest with slightly different temperature exposure, allowing ants to move between nests and behaviorally thermoregulate to some extent. Finally, as a desert arboreal ant, *C. rohweri* lives in an exceptionally variable thermal environment and microbes found within this species of ant guts might be expected to withstand change in temperature better than many insects due to a long history of natural selection on both the host and its microbes.

Beyond temperature, other factors almost certainly influence change in the *C. rohweri* microbiome across time and space. In particular, a portion of the effect of season could be due to a likely seasonal change in diet. As *Cephalotes* are largely herbivorous and potential scavengers of microscopic foods, the diet of *C. rohweri* is likely altered by seasonal changes in availability of pollen, nectar, and small windblown food items (spores, bacteria) (De Andrade and Urbani 1999). This shift in diet may have contributed to the dramatic increase in similarity within winter colonies, as foraging intensity, and potentially abundance and diversity of plant-based resources, decreases in the winter. More information on seasonal variation in *C. rohweri* diet is needed to evaluate whether this is the case.

As we show here, temperature should be considered among the multiple factors that influence ectotherm microbiome composition in nature. Ectotherm microbe-based thermal sensitivity—in both composition and abundance--may be an important, but poorly characterized, determinant of organismal response to climate change and fitness in the face of changing thermal regimes. A set of general predictions regarding ectotherm traits, emerging from a synthesis of studies such as this one, would enable researchers to predict whether microbe-based phenotypic plasticity will help or hinder animals in a changing climate.

## Acknowledgements

This project based upon work supported by the National Science Foundation Postdoctoral Research Fellowship in Biology under Grant No. 1811641. We would like to acknowledge the Center for Population Biology Seed Grant program, for funding the initial graduate student collaboration among MM, AZ, and AH. We would also like to acknowledge Pima County, AZ for permission to conduct the study.

## Author Contributions

MM wrote the manuscript, performed the lab work, and performed the data analysis. MM, RV, and SP designed the study, interpreted results, and contributed to revisions. MM, AH, ME, and AZ conducted fieldwork and designed the growth chamber study. MM, AH, ME, and AZ designed field sampling protocols.

